# Microscopy analysis of delayed co-transcriptional splicing in highly expressed thyroglobulin gene

**DOI:** 10.1101/2024.06.04.597307

**Authors:** Simon Ullrich, Iliya Nadelson, Stefan Krebs, Helmut Blum, Heinrich Leonhardt, Irina Solovei

## Abstract

Transcription of the majority of eukaryotic genes is accompanied by splicing, a process that depends on the assembly of the spliceosome on introns. The timing of spliceosome assembly varies significantly between introns, transcripts, genes and species. While quick co-transcriptional intron removal has been demonstrated for many mammalian genes, most splicing events do not occur immediately after intron synthesis. In this study, we utilized the highly expressed *Tg* gene, which forms exceptionally long transcription loops (Leidescher et al., 2022), providing a convenient model for studying splicing dynamics using advanced light microscopy. Our single-cell oligopainting-based analysis revealed a delay in splicing several tens of kilobases downstream of a transcribed intron, a finding further supported by standard cell population analyses. We speculate that this phenomenon is due to the abnormally high transcription rate of the *Tg* gene, which may lead to a localized deficiency in splicing factors and, consequently, delayed spliceosome assembly on thousands of nascent transcripts decorating the gene. Additionally, we found that, in contrast to short introns (<10 kb), long *Tg* intron (>50 kb) is spliced promptly, providing further support for the idea that intron length may modulate splicing speed.

## INTRODUCTION

Eukaryotic genes differ from prokaryotic genes by the presence of intragenic non-coding sequences, or introns, which play a crucial role in regulation of gene expression, gene evolution, and protein heterogeneity. This eukaryotic innovation, however, comes at the cost of a complex, energy-consuming, and tightly regulated process of intron splicing (Alpert et al., 2017; de Almeida and Carmo-Fonseca, 2008; Muller-McNicoll and Neugebauer, 2013). Despite significant progress in splicing research, the timing of splicing events remains a subject of ongoing investigation and debate (Eichenberger et al., 2023; Gordon et al., 2021; Han et al., 2011; Merens et al., 2024).

The process of splicing involves the identification of key genomic elements that define an intron - such as the splice donor, splice acceptor, and branchpoint - by the spliceosome complex. After an intron is transcribed, the spliceosome excises it, typically with RNA polymerase II positioned about 45 nucleotides downstream of the intron’s 3’ border (Oesterreich et al., 2016). While co-transcriptional intron removal has been demonstrated for many mammalian genes (Brugiolo et al., 2013), most splicing events do not occur immediately after the synthesis of an intron (Merens et al., 2024). Over the last decade, several deviations from the canonical co-transcriptional splicing model have been described.

One example is post-transcriptional splicing, where introns are removed in any order after pre-mRNA polyadenylation (Coté et al., 2024; Drexler et al., 2020; Gaidatzis et al., 2015; Zeng et al., 2022). Another widespread variation is intron retention, in which one or more specific introns are preserved in polyadenylated RNA to regulate the release of mRNA into the cytoplasm (Braunschweig et al., 2014; Jacob and Smith, 2017; Pimentel et al., 2016). Finally, canonical splicing can be supplemented with recursive splicing (Zhang et al., 2018), a process of intron removal via multiple rather than a single splice step, with apparently stochastic splice site selection (Duff et al., 2015; Sibley et al., 2015; Wan et al., 2021).

We recently demonstrated that highly expressed long genes expand from their harboring loci and form microscopically resolvable transcription loops (TLs) (Leidescher et al., 2022). This TL extension has been attributed to the dense decoration of the gene with thousands of polymerases with attached nascent ribonucleoprotein granules (nRNPs), which impart intrinsic stiffness to the gene axis. In essence, the extent of TL expansion is influenced by the size of the attached nRNPs, which become increasingly voluminous over long exons or long introns, linking TL expansion to splicing dynamics. One of the studied genes, the thyroglobulin (*Tg*) gene, is particularly highly upregulated, far surpassing the expression levels of other genes, including many housekeeping and tissue-specific genes (Leidescher et al., 2022; Ullrich et al., 2023). Notably, although the Tg gene is not very long (180 kb in mice), it forms exceptionally long TLs that extend several microns into the nucleoplasm (Fig. 1A).

**Fig. 1.**
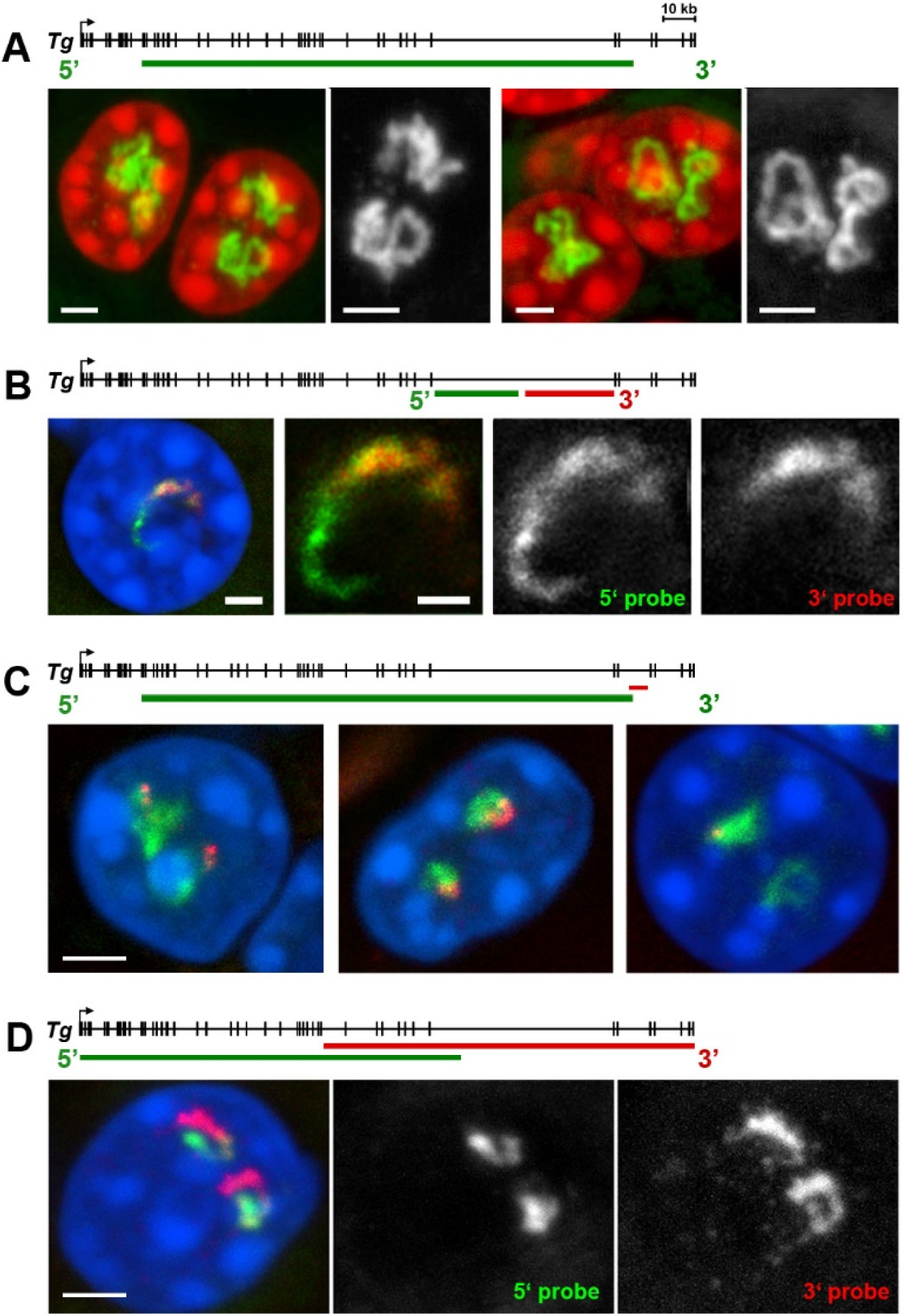
Strong extension of the *Tg* TLs allows transcription analysis using light microscopy. **A**, several examples of *Tg* TLs (*green*) within thyrocyte nuclei (*red*) detected by RNA-FISH using genomic BAC probe encompassing ca. 150 kb of the 180 kb long *Tg*. Note the volatile and strongly convoluted shape of TLs. **B,** labelling of intron 40 by RNA-FISH with oligoprobes for 5’ (*green*) and 3’ (*red*) halves of the intron. 5’ probe labels the entire intron, whereas the 3’ probe labels only half of the intron. The thickness gradient of the nRNAs increasing towards the 3’ end reflects the direction of transcription. **C,** the oligoprobe for the intron 44 at the 3’ end of the gene (*red*) in combination with BAC probe for the whole loop (*green*) after RNA-FISH. Three examples demonstrate similar intron signals (*left*), different sizes (*mid*) and signal on only one of the alleles (*right*). **D,** labelling of the *Tg* TLs with two BAC clones encompassing 5’ (*green*) and 3’ (*red*) halves of the loops by RNA-FISH. Although the probes overlap in the middle of the gene, on a global scale, they label the TLs sequentially as a result of co-transcriptional splicing. Every panel is accompanied by schematics (above images) illustrating the coverage of the gene with probes. Images are projections of short confocal stacks; nuclei are counterstained with DAPI. RGB panels are supplemented with grey scale images of the RNA signals. Scale bars: A,C,D, *2 µm*; B, *1 µm* and *0.5 µm* in close-ups.

The substantial extension of the *Tg* gene is somewhat counterintuitive, given its conventional organization: it contains 48 exons ranging from 66 to 233 nt and introns ranging from 197 to 9,375 nt, with only one large intron of approximately 54 kb in length. The *Tg* mRNA is only 8.4 kb long, and assuming quick co-transcriptional splicing, the lengths of nascent RNAs associated with *Tg* TLs should progressively decrease after each exon. This would result in smaller nRNPs decorating the gene, which should, in theory, permit the gene axis coiling and thus limit gene expansion. However, the *Tg* TLs expand more significantly compared to genes with long exons (e.g., *Ttn*) or long introns (e.g., *Cald1*) (Leidescher et al., 2022). One plausible explanation for the remarkable extension of *Tg* TLs) is a delay in intron splicing. Such a delay would result in long nRNAs containing not only exons but also multiple unspliced introns, leading to the formation of bulky nRNPs.

The *Tg* gene is exclusively expressed in thyrocytes, the secretory epithelial cells organized into closed follicles within the thyroid gland. When isolated from tissue and incubated *in vitro*, thyrocytes rapidly lose their identity and silence the *Tg* gene (Ullrich et al., 2023). Consequently, modern methods for studying splicing dynamics, such as live-cell imaging (Martin et al., 2013) or metabolic labeling of nRNAs (Schwalb et al., 2016), or POINT technology (Sousa-Luis et al., 2021), etc., cannot be applied in this context.

To test our hypothesis regarding delayed splicing of *Tg* nascent RNAs (nRNAs), we employed several methods compatible with thyroid tissue analysis. Standard cell population techniques, such as PCR and nanopore sequencing, did not provide sufficient information. Therefore, we conducted single-cell analyses using oligopainting to label consecutive *Tg* introns, followed by rigorous formal analysis of 3D images. Collectively, the data indicate a significant delay in the splicing of *Tg* introns, which we hypothetically attribute to the exhaustion of local splicing machinery caused by the gene’s high transcriptional activity.

## RESULTS & DISCUSSION

The remarkable length of *Tg* TLs (Fig. 1A) makes them as a unique and attractive model to study transcription and splicing, through light microscopy, as demonstrated by the following three examples. (I) RNA-FISH with differential labeling of the two halves of the 54 kb *Tg* intron enables us to observe the dynamics of intron splicing. Oligoprobes targeting the 3’ half label the second portion of the intron as anticipated, while oligoprobes for the 5’ half label the entire intron (Fig. 1B), suggesting that this long intron is likely excised as an entire piece. Admittedly, since the resolution of light microscopy does not allow for the analysis of individual nascent RNAs, we cannot completely rule out the possibility that some nascent transcripts undergo recursive splicing. However, the strong signal over the intron and gradient of snRNA suggest that, if recursive splicing does occur, it is not widespread. (II) RNA-FISH using oligoprobes targeting nRNA over 5 kb region of the last *Tg* intron reveals transcriptional bursting of the gene. Approximately equal numbers of nuclei displayed oligoprobe signals that were either similar in size on both alleles, different in size, or present on only one allele (Fig. 1C). The absence of RNA signals suggests that this intron, at the specific time of observation, remained in a transcriptional pause, while signals of differing sizes reflect nonsynchronous transcriptional bursting. (III) Our previous studies of TLs formed by long highly expressed genes have demonstrated co-transcriptional splicing but only on a large scale. RNA-FISH with two BAC probes, primarily targeting 5’ and 3’ introns, sequentially labels the TLs: 5’ introns are extensively spliced out as transcription proceeds toward the 3’ end of the gene, rendering them undetectable with the 5’ probe (Fig. 1D; see also (Leidescher et al., 2022; Ullrich et al., 2023)). Undeniably, due to the limited resolution of FISH with BAC probes, this experiment does not allow for a detailed assessment of splicing dynamics.

### RNA signals of *Tg* introns are disproportionally large

To study splicing dynamics at higher resolution, we labeled individual introns with oligoprobes using SABER-FISH (Kishi et al., 2019). The oligoprobes were designed to target the first 1–3 kb of each intron to enable labeling of the entire intron. We selected six sequential introns ranging in size from 2 to 7 kb, located at the 5’ end of the gene (Fig. 2A), and three introns at the 3’ end of the *Tg* gene, including the longest intron (40) with a length of 54 kb (Fig. 2B). Each intron was hybridized pairwise with a reference BAC probe in RNA-FISH. The BAC probe labeled nRNAs, including introns and exons, along a 153 kb region in the middle of the gene. Surprisingly, visual pairwise comparisons of the single intron signals with the BAC signal revealed disproportionately large signals for all introns except intron 40 (Fig. 2A,B; Fig. S1A,B).

**Fig. 2.**
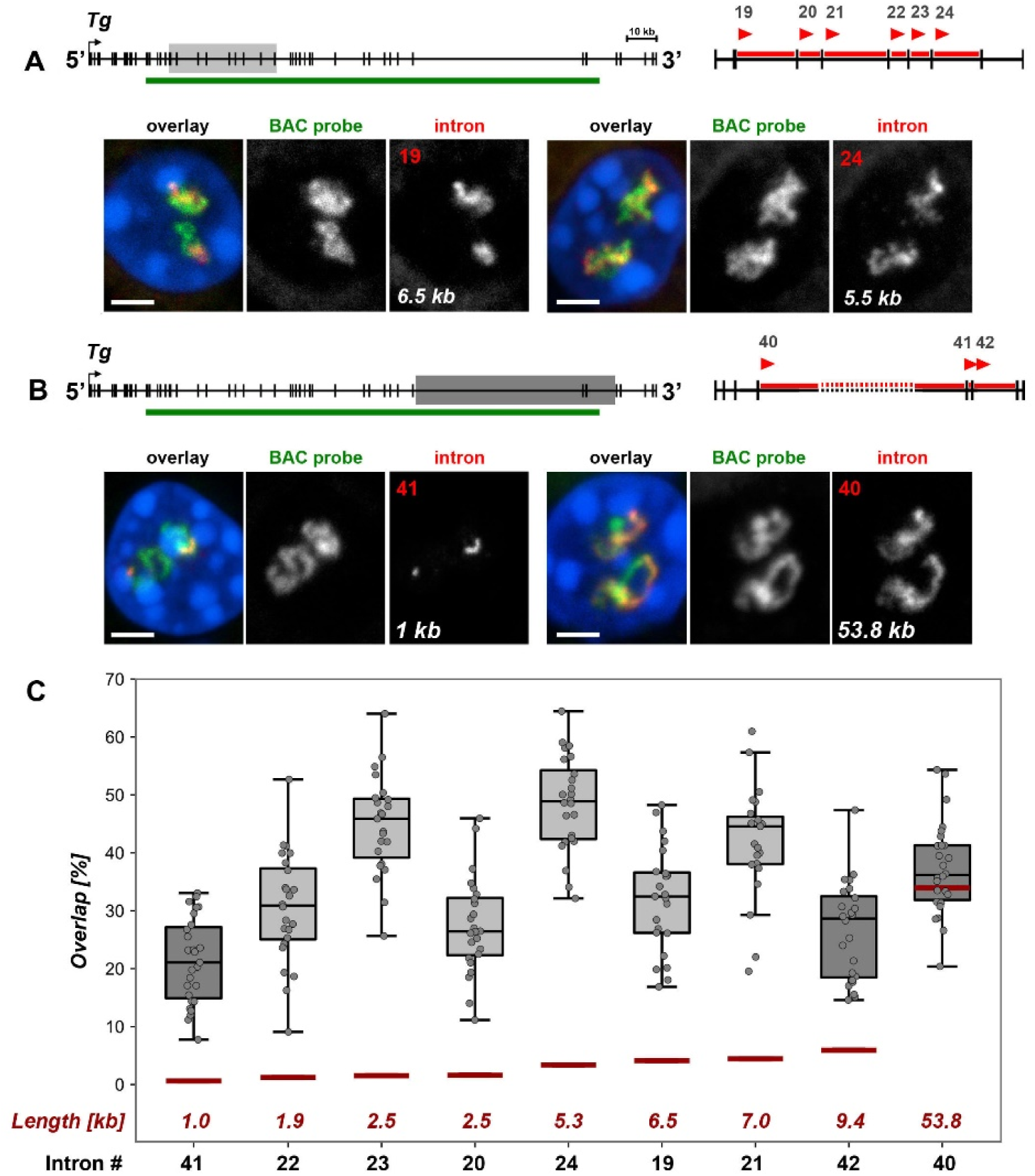
RNA signals of single *Tg* introns are disproportionally large in comparison to RNA signal of a genomic probe. **A, B,** pairwise comparison of RNA signals of single intron *versus* BAC probe (153 kb). In the gene schematics, sampled regions in the 5’ end (A) are marked with *light-grey rectangle* and in the 3’ end (B) - with the *dark-grey rectangle* gene regions. The green lines indicate coverage by the BAC probe; on the close-ups of the sampled regions, the *arrowheads* mark the 5’- end of introns, for which oligoprobes were designed. Exemplifying images of thyrocytes after RNA-FISH with pairwise probes (BAC, *green*; introns, *red*) are shown below the schematics. For clarity, RGB images are supplemented with grey scale images of RNA signals. Numbers and lengths of introns are indicated on the grey scale intron panels. Scale bars: *2 µm*. For more examples of nuclei in this experiment, see Fig. S1. **C,** boxplots showing the overlap ratio of intron signals with the BAC probe from both gene regions. Introns are arranged on the X-axis from smallest to largest; their numbers indicated below and their lengths - above the X-axis. Thick lines within boxes indicate the medians; boxes show 25^th^ and 75^th^ percentiles; whiskers represent the smallest and the largest values within 1.5 times the interquartile range; grey dots are individual data points. The number of evaluated nuclei ranged between 23 and 27. Note that the signal overlap of similarly sized introns can be significantly different, as it is the case for intron 20 and intron 23, both 2.5 kb in size (p < 0.001). On the contrary, introns that differ in their size, such as intron 23 (2.5 kb) and intron 24 (5.5 kb), can show similar overlap ratios (p > 0.05, one-way ANOVA with Tukey’s multiple comparisons post hoc). Expected overlap proportions based on the genomic size of the probes are marked with red lines. Note that only in case of the long intron 40, expected and measured overlaps are similar; for all the other introns, the measured overlap exceeded the expected value 6- to 10-fold.

To quantify signal disproportionality, we developed a dedicated workflow for 3D image analysis of confocal stacks. The program automatically segments and separates nuclei, identifies and segments RNA signals, corrects for chromatic shift between channels, and calculates the overlapping volume (V) between BAC and intron signals as the ratio Voverlap/(Vintron+V_BAC_) (for details, see the Methods). Because RNA signals from the two alleles within a single nucleus are often inseparable and may vary in size due to transcriptional bursting, the overlapping volume was calculated for both alleles together. This quantitative analysis confirmed the striking disproportionality of intron signals (Fig. 2C).

Based on the genomic sizes of the probed regions, the expected overlap proportion should correspond to the calculated intron/BAC length ratio, as indicated by the dark red lines at the bottom of the graph in Fig. 2C. However, the observed overlap was much higher ranging from 26% to 49%. For example, the expected overlap between intron 21 (7 kb) and the BAC (spanning 153 kb of the gene), is approximately 4%, but the measured overlap was over 40% (Fig. 2C). Interestingly, and somewhat counterintuitively, we observed a significantly larger overlap for short introns (1–10 kb), but not for the longest intron 40 with overlap value close to the expected ratio, i.e. around 35% (Fig. 2C).

The disproportionality of RNA intron signals suggests that a probe for an intron labels not only nRNAs transcribed from the targeted intron but also nRNAs from downstream regions of the gene. We hypothesized that this extra-labeling of the *Tg* TL could be explained by a delay in splicing along the gene body (Fig. 3A-D). Additionally, the observation that introns of different lengths exhibit similar overlap volumes may indicate distinct splicing dynamics for different introns.

**Fig. 3.**
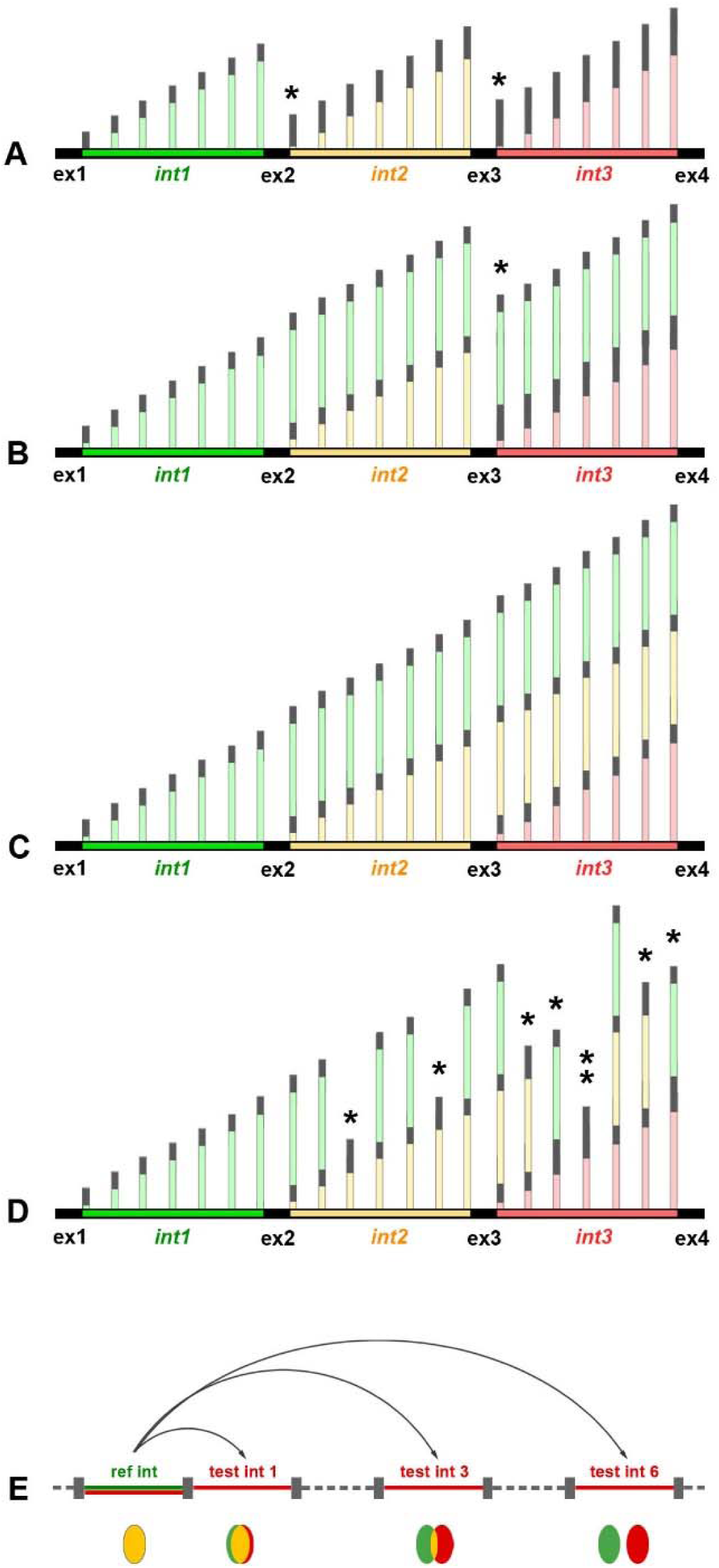
Schematics of possible splicing scenarios (A-D) and RNA-FISH experiments for estimation of splicing delay (E). Strictly co-transcriptional splicing (**A**) implies that introns are excised after polymerases cross the 5’ exon border. Singular (**B**) or consecutive (**C**) introns are excised synchronously on a certain distance from a transcription site. (**D**) Introns are excised in random nascent transcripts in a non-synchronous manner. Note that (B), (C) and (D) scenarios are not mutually exclusive. (E) Delay in splicing can be estimated by the degree of colocalization of RNA-FISH signals through comparison of a *reference*-intron (*green*) with *test-*introns (*red*). High signal colocalization (*yellow spot*) means delay in intron excision; absence of signal colocalization (separated *green* and *red spots*) indicates that the *reference*-intron is already excised from nRNAs decorating the *test-*intron. *ex*, exons; *int*, introns; *asterisks* mark spliced transcripts. Schematics of nRNAs (*vertical lines*) are not to scale to the gene regions (*horizontal lines*).

### Unspliced Tg introns are detectable by PCR

To confirm instances of splicing delay, we extracted total RNA from mouse thyroids and performed PCR using primers designed for regions spanning intron-exon-intron junctions, located approximately 250 bp upstream of an exon’s 5’ boundary and 750 bp downstream of its 3’ boundary. For all four tested regions, the intron-exon-intron sequences were successfully amplified, confirming that splicing of these introns is delayed by at least 0.75 kb (Fig. S2). This finding aligns with previous studies showing that, in some human genes, fully transcribed introns may persist within nascent RNAs for up to 2 kb downstream of the splice junction (Drexler et al., 2020; Sousa-Luis et al., 2021). However, the extent of *Tg* intron signal disproportionality observed in microscopy suggested that the splicing delay during *Tg* transcription is much greater than what is technically detectable *via* PCR. To address this, we sought to detect *Tg* nascent RNAs at a high-throughput level to examine longer fragments of nascent transcripts.

### Nanopore RNA sequencing demonstrates massive retardation of *Tg* introns in nRNAs

Detecting delayed splicing from bulk RNA-seq data is challenging, primarily due to the short length of RNA-seq reads, which are typically insufficient for characterizing single or multiple introns within individual transcripts. Recently, several highly efficient parallel methods for nascent RNA analysis have emerged, including GRO-seq (Core et al., 2008), NET-seq (Mayer and Churchman, 2016), TT-seq (Schwalb et al., 2016), and POINT technology (Sousa-Luis et al., 2021). These techniques require either metabolic labeling or the isolation of chromatin-associated RNA, which presents significant challenges for tissue-derived cells, particularly when the tissue is difficult to dissociate or when cultivating primary cells does not preserves their transcriptional program -both of which apply to the thyroid gland (Ullrich et al., 2023).

Although Nanopore sequencing allows long-read direct RNA sequencing of nascent transcripts (Garalde et al., 2018) and has been used as a powerful tool for dissecting the intricacies of RNA processing dynamics (Parker et al., 2020; Sousa-Luis et al., 2021), in the case of thyroid tissue, we are unable to draw reliable conclusions, which we tentatively attribute to the small amount of RNA we obtained. According to the manufacturer, to obtain accurate and reliable results with this method, the input RNA should be in the range of 1000 ng of total RNA. Given that thyrocytes comprise only 60% of the thyroid gland and that a single mouse thyroid yields less than 200 ng of total RNA, with only a small proportion representing nascent RNAs, the required quantity of thyroids and sacrificed mice for a single experiment would be unreasonably high. Therefore, we sought an alternative approach to estimate the magnitude of the splicing delay.

To elucidate the dynamics of delayed splicing events, we used an unconventional Nanopore sequencing workflow, enriching the sequencing library for nascent RNAs by depleting polyA RNA from the total RNA isolated from the tissue. Only intron-containing reads were included in the analysis; the length distribution of intron-containing Tg RNA reads indicated even sequencing coverage of nascent *Tg* RNAs (Fig. 4A, gray columns).

**Fig. 4.**
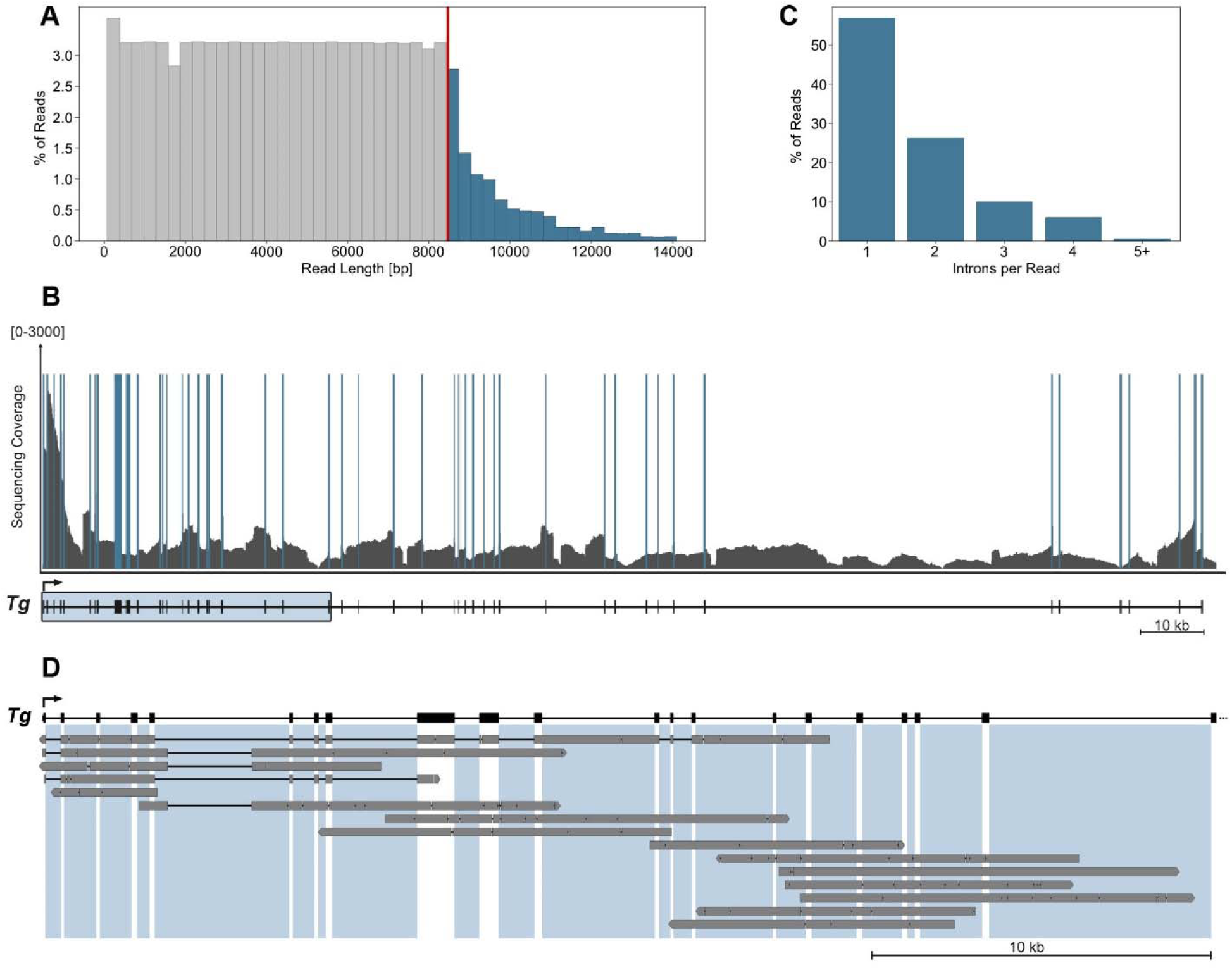
Nanopore sequencing confirms massive splicing delay of *Tg* introns. **A,** length distribution of nascent *Tg* RNA sequencing reads containing introns. X-axis represents the length of the individual sequencing reads. Each bar in the histogram corresponds to the frequency of reads with length intervals of 300 bp. Y-axis represents the number of reads. The red line designates the length of a fully spliced *Tg* mRNA (8462 bp). **B,** sequencing coverage of nascent RNAs along the Tg gene. *Solid blue bars* depict exons, *grey lines* depict introns. The *Tg* gene schematic is shown below the graph. The Y-axis is cut-off at 3000 reads to highlight the intron coverage. **C,** distribution of intron calls per intron containing read. X-axis represents the number of introns contained within individual sequencing reads. Each bar on the histogram represents the frequency of reads. In total, 10421 intron containing reads aligning to the *Tg* gene were assessed. **D,** graph exemplifying several individual read alignments mapped to the 5’ region of the *Tg* gene, outlined by the box in B. The *thick grey lines* represent exons and unspliced introns; the *thin black lines* represent spliced out regions. *Areas shaded with light-blue* highlight intronic regions within the gene sequence.

Multiple reads contained one or more fully transcribed introns, while others included only partial intron regions, making it difficult to distinguish between introns that had already been transcribed and those still in the process of transcription. The read lengths spanned up to 8.5 kb, which is consistent with the size of fully spliced and mature *Tg* mRNA. However, a fraction of reads exceeded 8.5 kb, reaching up to 15 kb (Fig. 4A, blue columns), clearly indicating the presence of unspliced introns. Coverage across all intronic regions within the *Tg* gene was comprehensive and uniform (Fig. 4B), highlighting the robustness of our approach in capturing nascent transcripts.

Unsurprisingly, the distribution analysis of intron-containing reads revealed a predominant presence of reads with only a single intron. However, a noticeable fraction of reads incorporated multiple introns, with some containing up to eight introns (Fig. 4C). Recent studies using techniques such as PRO-seq (Reimer et al., 2021) and POINT-seq (Sousa-Luis et al., 2021) have demonstrated that uncleaved transcripts can accumulate at the 3’ end of genes due to the presence of unspliced introns, thus contributing to the population of polyA(-) transcripts. Our analysis showed an even distribution of reads containing unspliced introns around the 3’ end of the *Tg* gene (Fig.4B). While we detected unspliced transcripts downstream of the *Tg* polyA site, they represented only a small fraction of the polyA(-) transcripts - around 350 reads out of approximately 110,000 total reads aligning to the last *Tg* exon - and therefore did not significantly contribute to the analyzed intron-containing fraction of nascent RNAs.

Examination of individual reads revealed instances where delayed splicing was clearly manifested, with multiple unspliced introns present in a single read, sometimes combined with already spliced introns (Fig. 4D). Our previous Nanopore sequencing data of the poly(A)-enriched mRNA fraction extracted from mouse thyroid tissue showed no intron retention in the polyA(+) *Tg* mRNA fraction (Ullrich et al., 2023). Therefore, we ruled out the possibility that the introns in our reads are due to regulatory intron retention for controlled mRNA export and concluded that our data reflect a splicing delay. Together, our findings demonstrate a delay in splicing across both single and multiple introns within individual *Tg* transcripts, however, due to technical limitations, we are unable to determine the exact duration of the splicing delay for individual introns.

### Pairwise comparison of intron signals confirms splicing delay

We reasoned that the considerable length of *Tg* TLs (Fig. 1A) makes it possible to examine splicing at the single-cell level using RNA-FISH and microscopy. First, since RNA-FISH does not require denaturation of DNA in tissue sections, probes targeting introns will not hybridize to the gene DNA, ensuring reliable detection of nRNAs. Second, visualizing differentially labeled introns in a pairwise manner would enable a signal comparison between a reference intron and downstream introns (Fig. 3E). We hypothesized that if there is a delay in splicing of an intron, its signal would colocalize with the signal from the subsequent intron or even multiple introns. Based on our observation of co-transcriptional splicing on the *Tg* gene at a global scale (Fig. 1D), we anticipated that intron signals would separate as the genomic distance between them increases (Fig. 3E).

For this analysis, we used the same regions at the beginning and end of the gene as in the previous RNA-FISH experiment (Fig. 2, S1). Within each region, we selected a reference intron at the 5’ end - specifically, introns 19 and 38 - and compared their signals with those of differentially labeled downstream test introns. Additionally, we included positive and negative controls for both regions. For the positive controls, which demonstrate maximally positive correlation, we used the same reference introns labeled with two distinct fluorophores. For the negative controls, we compared the reference intron with downstream introns positioned at a distance of no less than 50 kb (Fig. 5, S3).

**Fig. 5.**
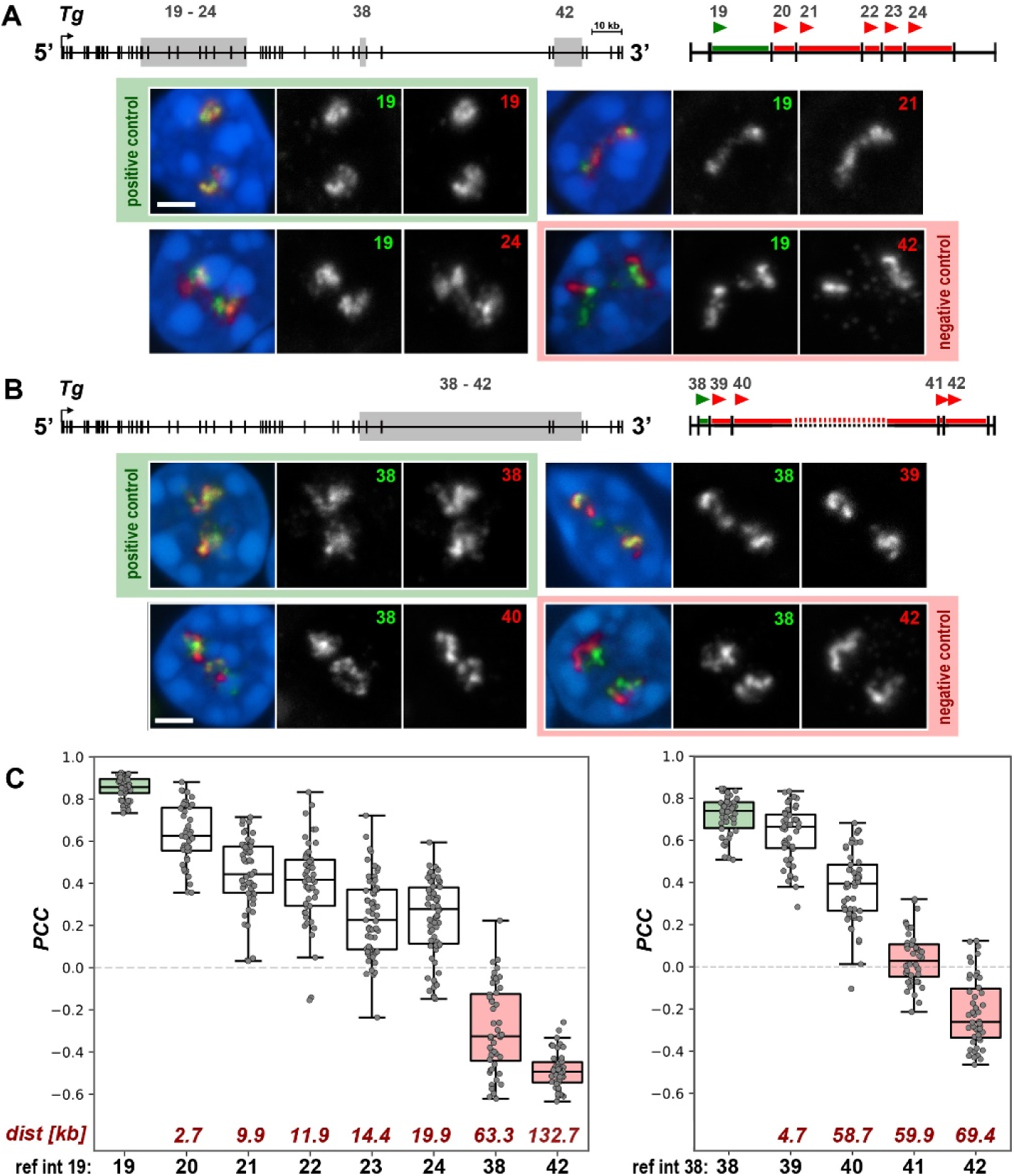
Pairwise comparison of *Tg* intron signals suggests splicing delay for tens of kb. **A, B,** pairwise comparison of RNA signals between single introns. Regions of intron sampling are marked on gene schematics by *grey rectangles*. On the close-ups of the sampling regions, *green lines* and *red lines* mark the reference- and test-introns, respectively; *arrowheads* mark 5’-end of introns, for which oligoprobes were designed. Exemplifying images of thyrocytes after RNA-FISH with pairwise intron probes are shown below the schematics. The reference-introns 19 (A) and 38 (B) are *green*; the test-introns are *red*. For clarity, the overlaid RGB images of RNA signals are supplemented with grey scale images. Numbers of introns are indicated on the grey scale panels. Images are projections of short confocal stacks. Scale bars: *2 µm*. For more examples of nuclei in this experiment, see Fig. S3. **C,** boxplots showing the PCC values for pairwise intron comparison for regions on A (*left*) and B (*right*). Introns are arranged along the X-axis according to their genomic positions. Numbers below the axis are intron numbers; numbers above axis are distances of test-introns from the reference-intron. Thick lines within boxes indicate the medians; boxes show 25^th^ and 75^th^ percentiles; whiskers represent the smallest and the largest values within 1.5 times the interquartile range; grey dots are individual data points. Number of evaluated nuclei ranged between 40 and 50. Positive and negative controls are highlighted with *light-green* and *light-red* colors, respectively.

For each comparison, we estimated the colocalization of introns using the Pearson’s correlation coefficient (PCC). Differentially labeled nRNAs of the reference and test introns were detected in two spectrally separated channels, enabling pixel intensity-based colocalization analysis. This analysis involves evaluating the intensity of each pixel in one channel against the intensity of the corresponding pixel in the other channel, generating a correlation coefficient (Cordelieres and Bolte, 2014; Cordelières and Zhang, 2020). For this purpose, we modified our analysis pipeline to calculate the PCC between signals in two channels within a 3D volume (for details, see Methods). Importantly, the collected image stacks were corrected for both axial and lateral chromatic shifts between the spectrally separated channels. To accomplish this, stacks of 0.5 µm beads labeled with four fluorophores (TetraSpeck™ Microspheres) were collected during each acquisition session and processed alongside the samples.

As expected, the signals of both fluorophores in positive controls visually overlap (green fields in Fig. 5A,B) and the mean PCCs for them were close to 1 (Fig. 5C). In contrast, the signals for the negative controls were visually separated (pink fields in Fig. 5A,B), and their PCC values were negative (Fig. 5C). It is important to note that the anti-correlation does not carry any specific biological meaning but instead indicates signal separation. The mean PCC values for seven pairwise comparisons between the reference and tested introns ranged between those for the positive and negative controls, consistently decreasing as the genomic distances between them increased (Fig. 5C), which well corresponded to the visual estimates (Fig. 5A,B; Fig. S3A,B).

Our microscopy analysis tentatively suggests that intron splicing may be delayed over tens of kilobases, i.e., assuming an average polymerase speed of approximately 3.8 kb/min (Singh and Padgett, 2009), for up to tens of minutes after transcription. Undeniably, however, our microscopy approach has its limitations. RNA-FISH detects hundreds of nascent transcripts densely arranged along the *Tg* TLs, making it impossible to identify a single nascent transcript within this mass. Consequently, we cannot determine whether splicing is delayed in all nascent RNAs containing a particular intron. However, based on the bright intron signals, this delay appears to be present in the majority of them.

Additionally, we would like to point out that our analysis involves structures smaller than 0.5 - 1 µm, approaching the resolution limit of light microscopy. As a result, we cannot completely rule out the possibility that the colocalization of intron signals is influenced, to some degree, by the coiling of the gene axis, which could bring neighboring introns into close proximity and mix their signals. However, our observations of the disproportionality of single intron signals compared to the nearly full-length TL (Fig. 2) suggested that gene axis coiling alone cannot fully explain the phenomenon we observed in the pairwise intron comparison.

### Splicing delay is not observed in moderately expressed gene

The Tg gene is characterized by an exceptionally high expression level of approximately 23,000 TPM (Leidescher et al., 2022). We hypothesize that this high transcription level may lead to a localized depletion of the splicing machinery, preventing timely spliceosome assembly and thereby causing splicing delays. Unfortunately, we are unable to test this hypothesis directly by manipulating *Tg* expression in a mouse model due to ethical constraints and the fact that transcription inhibition results in TLs withdrawal (Leidescher et al., 2022), which would prevent microscopy analysis. Additionally, we cannot analyze other genes with similar length and high expression levels, as we are unaware of any other mammalian genes that meet these criteria.

We reasoned that, as an indirect test, we could examine a gene with a length comparable to that of *Tg* but with lower expression levels to assess whether it also exhibits splicing delay. For this, we selected the *Cald1* gene, which is 177 kb long and expressed in cultured myoblasts at approximately 1,800 TPM. The expression level of *Cald1* is 13-fold lower than that of *Tg* and can be considered moderate. In line with this lower transcription level, the gene forms small, though microscopically resolvable, transcription loops (Leidescher et al., 2022).

Similar to the *Tg* gene, we tested the splicing dynamics of *Cald1* through pairwise colocalization comparison of intron signals (Fig. 6). The first *Cald1* intron was used as a reference, and colocalization of its signal was measured pairwise with the four subsequent test introns (Fig. 6A). Visual inspection of confocal stacks showed that only the signals from the reference intron labeled with two fluorophores exhibited colocalization. The signals for the subsequent test introns, despite small size of the Cald1 TLs, were evidently distinguishable though from the reference (Fig. 6B). In line with these observations, the PCC values between the reference and test introns were at zero or below (Fig. 6C). Taken together, our data indirectly support the hypothesis that local splicing dynamics may be influenced by transcription levels.

**Fig. 6.**
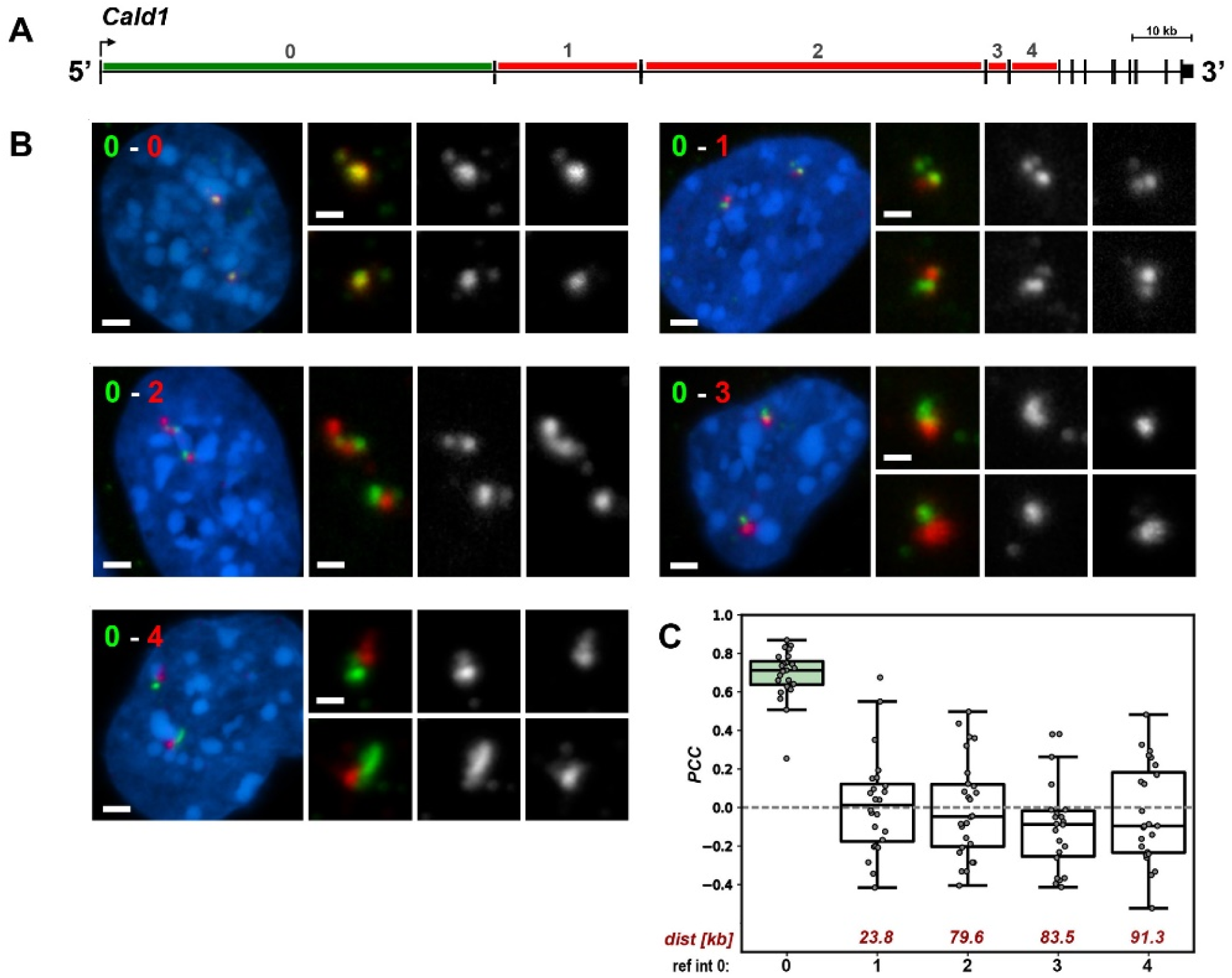
Splicing delay is not detected in the moderately expressed *Cald1* gene. **A,** schematics of the Cald1 gene with marked reference intron (*green line)* and test-introns (*red lines*). Intron numbers are indicated above the gene axis. **B,** exemplifying images of myoblasts after RNA-FISH with pairwise intron probes. The reference-intron 0 is *green,* the test-introns 1-4 are *red*. Numbers of compared introns are indicated in the upper left corners. Close-ups of the signals are shown on the right and supplemented by grey scale images. Images are projections of short confocal stacks. Scale bars in panels with nuclei: *2 µm*, in close-ups: *1 µm*. **C,** boxplots showing the PCC values for pairwise intron comparison with introns arranged along the X-axis according to their genomic positions. Numbers below the axis are intron numbers; numbers above the axis are genomic distances of test-introns from the reference-intron. Thick lines within boxes indicate the medians; boxes show 25^th^ and 75^th^ percentiles; whiskers represent the smallest and the largest values within 1.5 times the interquartile range; grey dots are individual data points. Number of evaluated nuclei ranged between 22 and 26.

### Splicing dynamics for short and long introns is different

The proportional size of the *Tg* intron 40 signal compared to the BAC signal (Fig. 2B,C) suggested that there is no significant delay in the splicing of this particular intron. To investigate this further, we labeled three sequential introns - 40 (53.8 kb), 41 (1 kb), and 42 (9.4 kb) - with three distinct fluorophores (Fig. 7A1). This setup allowed for three pairwise colocalization comparisons of the introns (Fig. 7A2). Consistent with previous experiments indicating splicing delay in *Tg* introns (Fig. 5), the two sequential introns, 41 and 42, showed a high degree of colocalization. Interestingly, the colocalization between introns 40 and 41 was significantly lower and even became negative between introns 40 and 42 (Fig. 7A3). For comparison, we used three shorter sequential introns, 22 (1.9 kb), 23 (2.5 kb), and 24 (5.3 kb) (Fig. 7B1). As anticipated, colocalization of the three differentially labeled introns resulted in high PCCs for all pairwise comparisons (Fig. 7B2, B3).

**Fig. 7.**
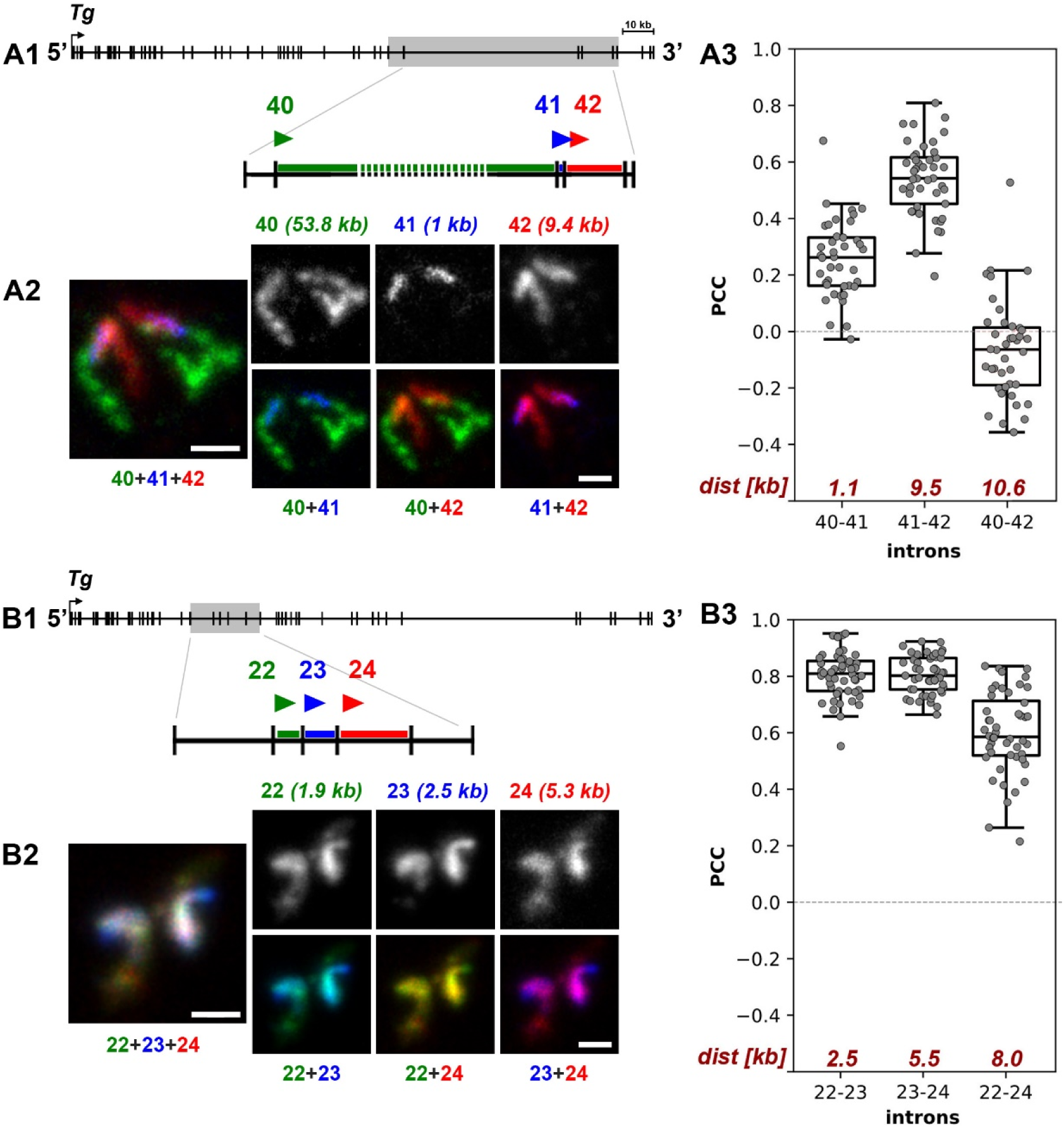
Long introns exhibit only small delay in splicing. Estimation of colocalization between three consecutive introns that include one long intron (*53.8 kb*) followed by two short ones (*1 and 9.4 kb*) (**A**) and between three consecutive short introns (*1.9, 2.5 and 5.3 kb*) (**B**). A1 and B1 are schematics of the gene with highlighted regions of the tested introns also showing the color code. A2 and B2 represent telling examples of intron colocalization. In addition to RGB pairwise overlays and an overlay of all three channels, grey-scale images of single introns are shown. Images are projections of several optical sections. Scale bars: *1 µm*. Boxplots in A3 and B3 show the PCC values for pairwise intron comparison. Numbers below the X-axis show intron numbers of compared introns; numbers above X-axis indicate distances between them. Thick lines within boxes indicate the medians; boxes show 25^th^ and 75^th^ percentiles; whiskers represent the smallest and the largest values within 1.5 times the interquartile range; grey dots are individual data points. Number of evaluated nuclei ranged between 40 and 50. Note, that by both visual inspection of the RNA-FISH signals and PCC measurement, the longest *Tg* intron 40 shows little correlation with the following small intron 41 and no correlation with intron 42, whereas both short introns 41 and 42 colocalize strongly (A), similarly to the three short introns in a different gene region (B).

The negative correlation between introns 40 and 42, which are separated by only 10.4 kb, contrasts with the data on the colocalization of introns 19-21 and 19-22: these introns are also separated by 10-12 kb but exhibit a PCC of around 0.5 (Fig. 5C). The prompt splicing of the longest *Tg* intron 40 suggests that intron length could indeed influence splicing speed. This observation aligns with a recent finding using POINT-nano technology, which demonstrated a positive correlation between intron size and splicing efficiency (Sousa-Luis et al., 2021).

We propose several hypothetical scenarios to explain this phenomenon. First, it’s possible that this intron undergoes recursive splicing, requiring less time for complete excision. Although this scenario is plausible, it likely applies only to a minority of transcripts, as we observe the entire intron labeling with the 5’ probe and steadily increasing gradient of nRNAs toward its 3’ end (Fig. 2B). The second scenario considers the pronounced extension of the intron 40 from the main *Tg* transcriptional loop (Fig. 2B; Fig. S1B;(Leidescher et al., 2022), positioning the intron in an environment with less competition for splicing machinery. Lastly, assuming that the RNAPII transcription speed is around 4 kb/min (Singh and Padgett, 2009), transcribing the entire 53.8 kb intron would take over 14 min, providing more time to assemble the necessary splicing machinery compared to shorter introns, which are transcribed in seconds to several minutes.

## CONCLUSIONS

Our microscopy analysis suggests that during transcription of the highly and perpetually upregulated *Tg* gene, splicing of small introns (<10 kb) is delayed over tens of kilobases, a finding confirmed by standard cell population analyses, including PCR and Nanopore sequencing. Intriguingly, the long *Tg* intron (>50 kb) stands out as an exception to this phenomenon, showing no measurable splicing delay.

We acknowledge that obtaining a precise estimation of the splicing delay for specific introns within individual transcripts by microscopy is not feasible. First, RNA-FISH generates massive signals from hundreds of nascent RNA transcripts densely decorating *Tg* subregions, rendering individual transcripts indistinguishable. Second, the limited resolution of light microscopy, even with super-resolution techniques (Leidescher et al., 2022), prevents detailed resolution of intron folding and its structure within the *Tg* gene. We tentatively attribute the splicing delay to the high transcriptional activity of the *Tg* gene, which may cause a localized depletion of the splicing machinery across multiple nascent RNA transcripts. As indirect supporting evidence for this hypothesis, we demonstrated that the introns of the *Cald1* gene, a gene of comparable length but with more than tenfold lower expression, are excised without a noticeable delay.

Splicing delay may follow different scenarios: (i) only certain introns experience delayed splicing (Fig. 3B); (ii) intron sequences persist across several exons before undergoing extensive splicing at a specific time point (Fig. 3C); (iii) in a subset of nascent RNAs, introns undergo asynchronous splicing downstream of their transcription sites (Fig. 3D). It is likely that a combination of these scenarios coexists during intense transcription. However, as noted above, the resolution of both Nanopore sequencing and microscopy analysis does not allow us to differentiate between these potential scenarios. For example, we could distinguish between cases illustrated by in Fig. 3A from Fig. 3B or Fig. 3C,D, but not between Fig. 3C and 3D, because RNA-FISH would yield similar signals in these two case.

Irrespective of the possible scenarios discussed above, delayed splicing provides an explanation for why *Tg* transcription loops extend so profoundly into the nuclear interior. The accumulation of introns in nascent RNAs gradually increases the size of nRNPs over tens of kilobases. Consequently, the increased bulkiness of these nRNPs enhances the stiffness of the gene axis, forcing the gene to extend further into the nuclear space - a phenomenon previously demonstrated in biological experiments involving splicing inhibition and computer polymer simulations (Leidescher et al., 2022).

Altogether, our findings illuminate the intricate and dynamic nature of RNA processing, highlighting underlying complexities in the regulation of splicing events during transcript maturation. Despite technical challenges and certain limitations, we demonstrated that working with tissue samples can be highly rewarding, as it often reveals *in vivo* phenomena and processes that may not be present in cultured cells *in vitro*. Additionally, our work underscores the power of microscopy and image analysis in transcription studies.

## MATERIAL AND METHODS

### Tissue collection and cryosections

Thyroid tissue sampling was executed in accordance with the European Union (EU) directive 2010/63/EU on the protection of animals used for scientific purposes and in compliance with regulations by the respective local Animal Welfare Committees (LMU; Committee on Animal Health and Care of the local governmental body of the state of Upper Bavaria; Germany). Mice were sacrificed by cervical dislocation after IsoFlo (Isofluran, Abbott) narcosis. Freshly dissected tissues were washed with PBS and then fixed with 4% paraformaldehyde (Carl Roth) solution in PBS for 12-20 h. After fixation, thyroids were washed with PBS, cryoprotected in a series of sucrose, and embedded in Tissue-Tek O.C.T. compound freezing medium (Sakura). Blocks were stored at - 80°C before cutting into 16-20 µm sections using a cryostat (Leica CM3050S). Cryosections were collected on Superfrost Plus slides (Thermo Scientific) and stored at - 80°C before use.

### Cell culture

The mouse myoblast cell line Pmi28 was grown in Nutrient Mixture F-10 Ham supplemented with 20% fetal bovine serum (FBS) and 1% Penicillin/Streptomycin at 37°C and 5% CO_2_. Cells were sub-cloned on coverslips pretreated with polylysine. After brief wash with pre-warmed PBS, cells were fixed with 4% paraformaldehyde (Carl Roth) solution in PBS for 10 min, washed with PBS (10 min x 3), permeabilized with 0.5% Triton X100 in PBS, washed with PBS supplemented with 0.01% Tween 20 (10 min x 3), equilibrated in 2xSSC and stored in 50% formamide in 2xSSC buffer.

### Probes

BAC clones encompassing the *Tg* gene (RP24-229C15, RP23-193A18, RP23-266I10) were selected using the UCSC genome browser and purchased from BACPAC Resources (Oakland children’s hospital) as agar stabs (https://bacpacresources.org/). BACs were purified via standard alkaline lysis or the NucleoBond Xtra Midi Kit (Macherey-Nagel), followed by amplification with the GenomiPhi Kit (GE Healthcare) according to the manufacturer’s instructions. Amplified BAC DNA was labeled with fluorophores using homemade conjugated fluorophore-dUTPs by nick translation (Cremer et al., 2008). Labeled BAC DNA was ethanol precipitated with 10-fold excess of Cot-1 (1 mg/ml; Invitrogen, 18440-016) and 50-fold excess of salmon sperm DNA (5 µg/µl; Sigma); pellet was dried in a SpeedVak, and dissolved in hybridization mixture containing 50% formamide, 1xSSC and 10% of dextran sulfate.

Genomic coordinates of the studied introns are listed in Supplementary Table 1. Oligoprobes for introns were generated using SABER-FISH protocol and described in detail previously (Leidescher et al., 2022). Briefly, oligos targeting intron sequences were designed using Paintshop (Hershberg et al., 2021) and ordered as oligo pools from Integrated DNA Technologies. The oligos were remapped if necessary for multi-colour imaging and extended to ∼500 nt using primer exchange reaction (Kishi et al., 2019). Finally, the probes were purified using PCR clean-up columns (Macherey-Nagel).

### FISH

RNA-FISH using BACs was performed on cryosections and cultured myoblasts as previously described. Denatured probes were loaded on sections under small glass chambers or on coverslips with myoblasts and sealed with rubber cement (for detail, see (Eberhart et al., 2012; Solovei and Cremer, 2010). Denaturation of sections and cells was omitted. Hybridizations were carried out in a water bath at 37°C for 1-2 days. After hybridization, rubber cement and chambers were removed, slides with sections or coverslips with myoblasts were washed with 2xSSC at 37°C, 3 x 30 min, and then with 0.1xSSC at 60°C 1 x 7 min. Hybridized SABER probes were detected by incubation with 1 μM fluorescently labeled detection oligonucleotides in PBS for 1 h at 37°C followed by washing with PBS for 10 min.

### Microscopy

Confocal image stacks were acquired using a TCS SP5 confocal microscope (Leica) using a Plan Apo 63/1.4 NA oil immersion objective and the Leica Application Suite Advanced Fluorescence (LAS AF) Software (Leica). Z step size was adjusted to an axial chromatic shift and typically was either 200 nm or 300 nm; XY pixel size varied from 20 to 60 nm. Chromatic shift was measured using Tetraspeck beads (0.5 µm).

### Image Analysis

All image sets (i.e., z-stack imaged in three spectrally separated channels) for an overlap or correlation experiment were processed automatically with custom python (v3.8.0; Python Software Foundation) scripts, making use of several packages: numpy (Harris et al., 2020), pandas (McKinney, 2011), skimage (Van der Walt et al., 2014), scipy (Virtanen et al., 2020), matplotlib (Hunter, 2007), seaborn (Waskom, 2021), tifffile (Gohlke) and napari (Ahlers et al.). The scripts are available upon request. The main steps are summarized below.

### Calculation of overlapping of FISH signals in 3D

First, each image set was corrected for chromatic shift between the channels. Next, the DAPI signal was binarised using Otsu’s method (Otsu, 1975) and cleaned from small artefacts. Subsequently, a watershedding procedure was employed to separate the nucleus in the centre of the image from neighbouring nuclei. Objects touching the image borders were removed and the object with the biggest volume was retained as the segmented nucleus.

Within the segmented nucleus, the signal of each FISH channel was segmented. First, the volume around each FISH signal was reduced by calculating and thresholding (Otsu) a local entropy image. The resulting mask was applied to the original FISH image, which was then also thresholded using Otsu’s method. The number of remaining objects was evaluated and a maximum of two objects (biggest volumes) per FISH channel was retained.

Finally, the overlap between objects in both FISH channels was calculated and compared to the total volume of objects in both FISH channels. Specifically, the signals of single introns were overlapped with a BAC probe delineating most of the gene. To ensure only results from correctly segmented signals remained, image sets where less than 80% of the segmented intron overlapped with the segmented BAC were excluded from the analysis.

### Calculation of correlation between FISH Signals in 3D

For the pixel-based colocalization analysis of FISH signals, images were processed as described above until the FISH segmentation. The volume around the FISH signals was further confined by summing both FISH channels and thresholding the result (Otsu’s method) while applying the nuclear mask. This reduced the influence of background fluorescence on colocalization calculations. The resulting mask was cleaned-up from artefacts and objects overlapping with the nuclear border were removed. The FISH signals were smoothed (Gaussian filter, σ=1) before the mask was applied to reduce the value differences between neighbouring pixels. Finally, the pixel-wise Pearson Correlation Coefficient (PCC) between the resulting volumes of both FISH signals was computed. The resulting correlation values were saved in .csv format and plotted using R Statistical Software.

### PCR analysis

Dissected thyroids were immediately placed in Lysis Buffer RA1 and homogenized using an ultra-turrax dispersing tool. Total RNA was isolated using the NucleoSpin RNA Kit (Macherey-Nagel). 1 μg of total RNA was reverse transcribed using Maxima H Minus Reverse Transcriptase (Thermo Scientific) with gene specific primers. PCR was performed in technical and biological triplicates using Phusion Plus polymerase (Thermo Scientific) and subsequently analysed on 1% agarose gel. All steps were performed according to the manufacturer’s recommendation. Distances between introns were calculated from the 3’ end of a reference intron to the 3’ end of a test intron. All used primers can be found in Supplementary Table 2.

### Nanopore sequencing

Freshly dissected thyroids were immediately placed in ice cold TriZol and homogenized using an ultra-turrax dispersing tool. The total RNA pool was depleted of Poly(A)+ transcripts using magnetic oligoT-beads (Lexogen). The sequencing library from the remaining RNA was generated using the PCR-cDNA Sequencing Kit (PCB111.24, Oxford Nanopore) and sequenced on a PromethION P24 on a R9.4.1 flowcell. The sequencing data were basecalled using Guppy v6.4.6 and mapped to the mouse genome (mm10) using minimap2 (Li, 2018).

## DATA ACCESS

All relevant data can be found within the article and its supplementary information. Data on Nanopore sequencing are deposited to SRA database with PRJNA1112134 accession number.

## COMPETING INTERESTS

The authors declare no competing or financial interests.

## ACKNOWLEDGEMENTS

We are grateful to Maria Carmo-Fonseca (Instituto de Medicina Molecular João Lobo Antunes, Faculdade de Medicina da Universidade de Lisboa) and Daniel Larson (Receptor Biology and Gene Expression, NIH) for fruitful discussions.

## FUNDING

This work has been supported by the Deutsche Forschungsgemeinschaft grants (SP2202/SO1054/2, project # 422388934 to IS, SPP 2202/LE721/17-1, project # 422857584 to HL, and SFB1064, project # 213249687 to HL and IS)

## AUTHOR CONTRIBUTIONS

*Conceptualization:* I.S.; *Methodology:* S.U., I.N., I.S.; *Validation:* S.U., I.N., I.S.; *Formal analysis:* S.U., I.N.; *Investigation*. S.U., I.N., I.S., S.K., H.B.; *Resources:* S.K., H.B.; *Data curation:* I.S., H.L.; *Writing - original draft:* S.U., I.S., I.N.; *Writing - review & editing*: S.U., I.S., I.N., H.L.; *Visualization*: S.U., I.S., I.N.; *Supervision:* I.S.; *Project administration*: I.S.; *Funding acquisition:* I.S., H.L.

---

**Fig. S1.**
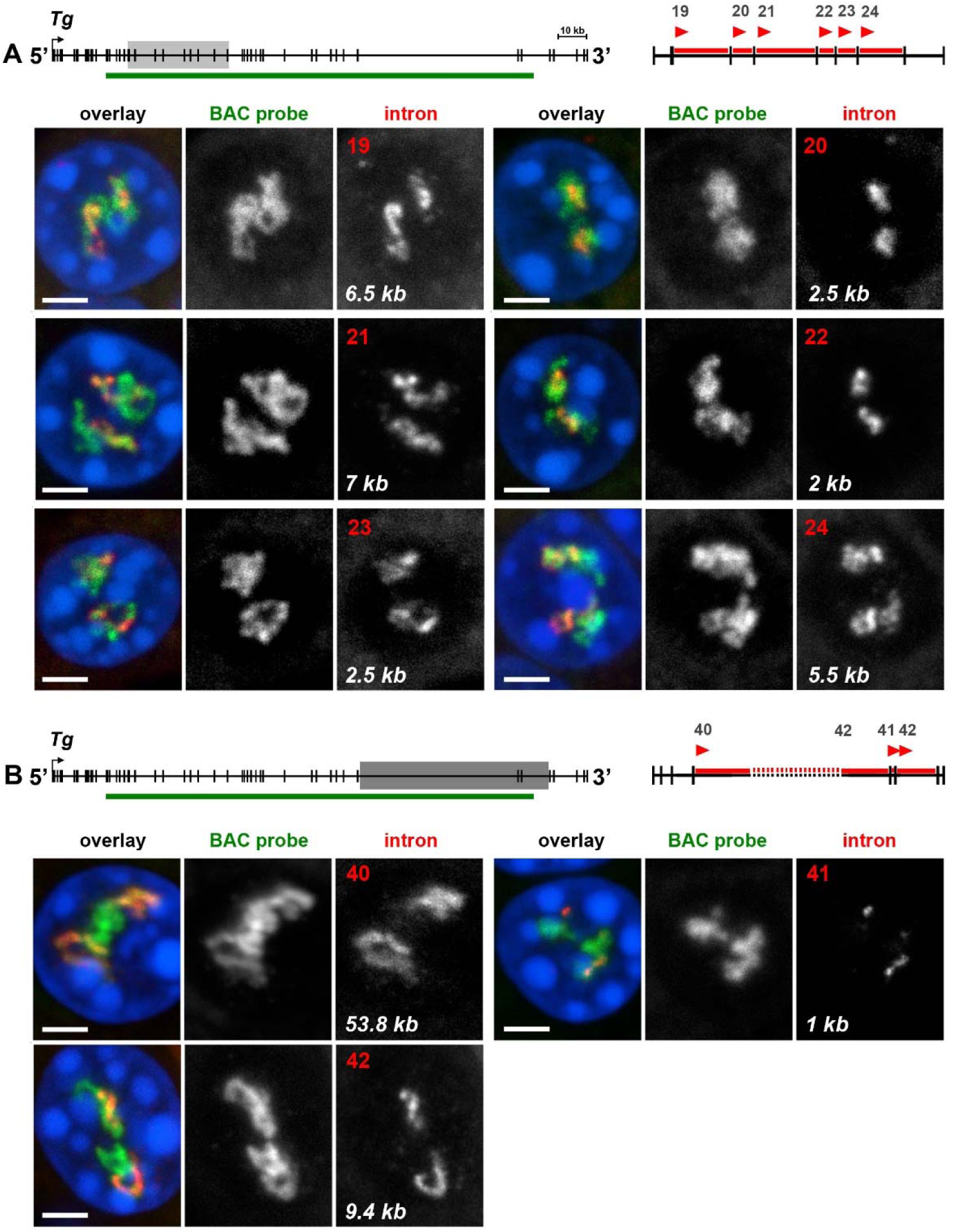
Overlapping between RNA signals of the *Tg* introns and BAC probe encompassing the middle of the gene (supplementary to Fig. 2). Pairwise comparison of RNA signals of a single intron *versus* BAC probe (153 kb) in the 5’ (**A**) and 3’ (**B**) gene regions. The regions are marked on gene schematics with *light-grey* and *dark-grey rectangles*, respectively. The *green lines* below gene schematics indicate localization of the BAC probe; the red arrowheads mark 5’-end of introns within the indicated regions, for which oligoprobes were designed. Images are projections of confocal stacks through thyrocytes after RNA-FISH with pairwise probes (BAC, *green*; introns, *red*). For clarity, RGB images are appended with grey scale images of RNA signals. Numbers and lengths of introns are indicated on the grey scale intron panels. Scale bars: *2 µm*.

**Fig. S2.**
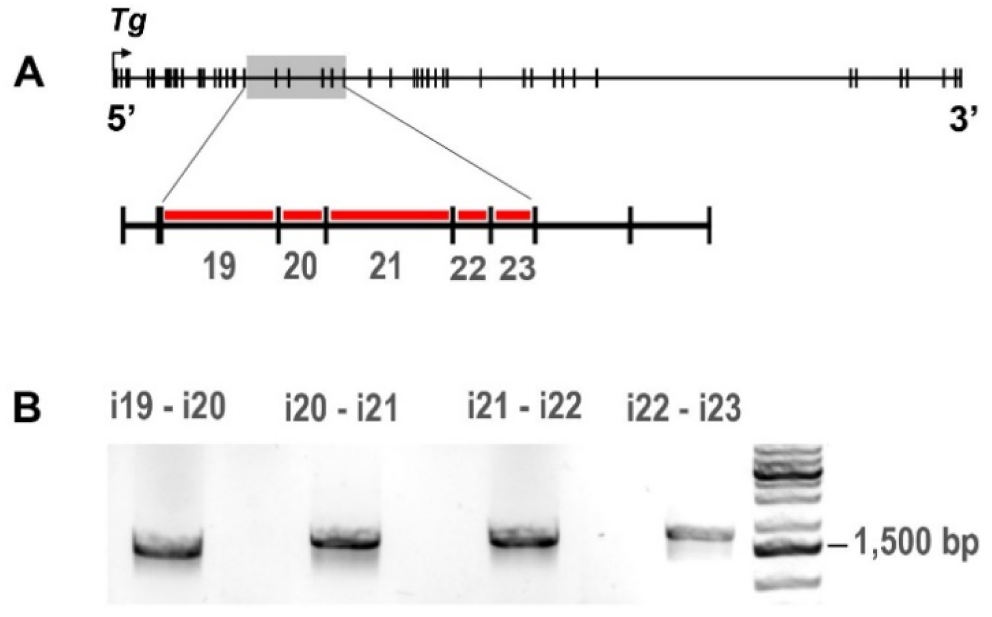
Unspliced *Tg* introns are detectable by PCR. **A**, Gene schematics showing localization of the tested *intron-exon-intron* sequences. In the enlarged view, the used introns are marked with red lines. **B**, PCR with primers designed from both sides of the tested exons: ca. 250 bp upstream of an exon 5’ border and ca. 750 bp downstream of its 3’ border. For all four tested exons, the *intron-exon-intron* sequences were successfully amplified.

**Fig. S3.**
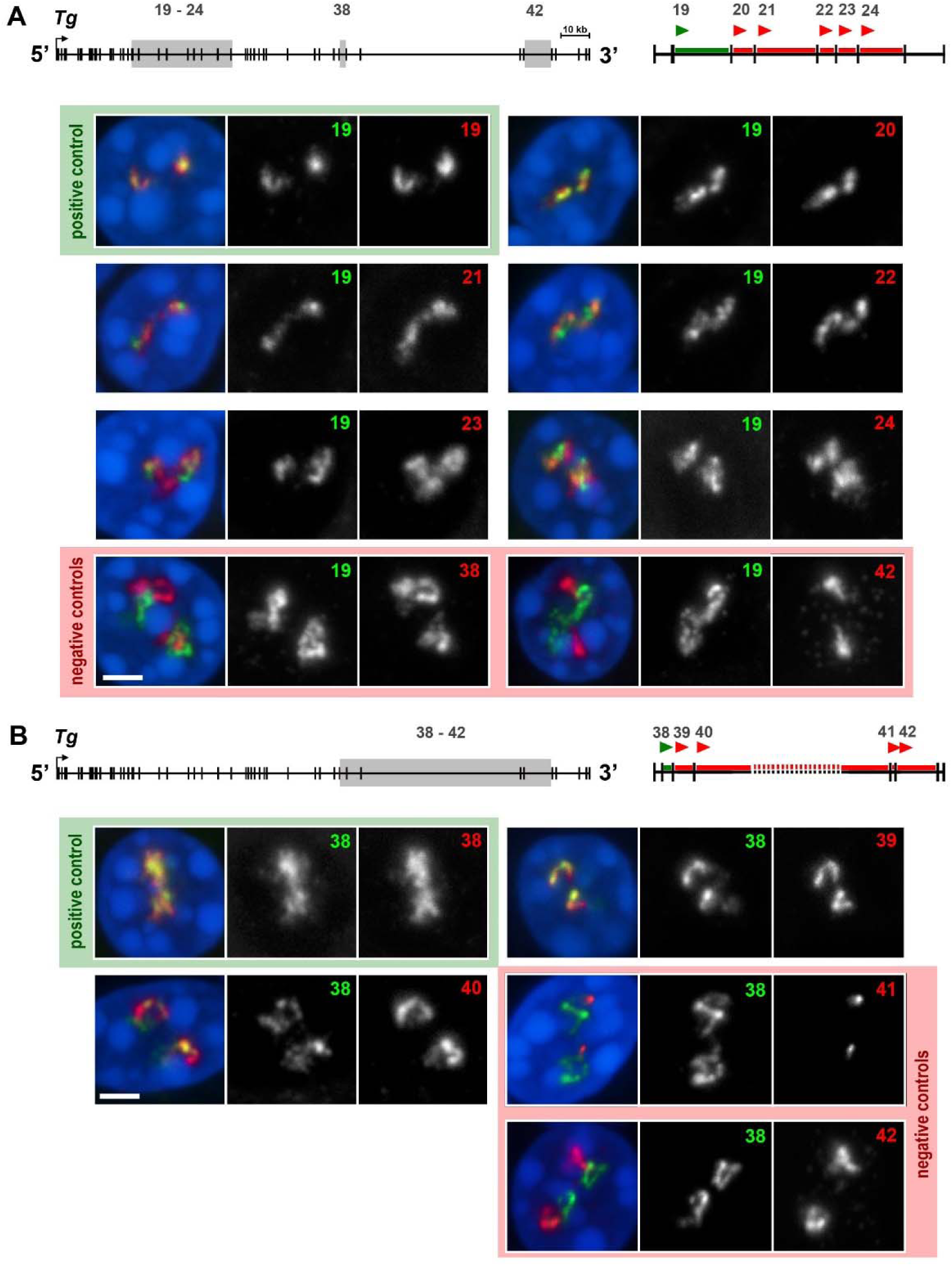
Pairwise comparison of *Tg* intron signals (supplementary to Fig. 5). **A, B,** exemplifying images of thyrocytes after RNA-FISH with pairwise comparison of RNA signals between single introns. Two regions with detected introns at the 5’ end (A) and 3’ end (B) are marked on gene schematics by *grey rectangles*. On the close-ups of the sampling regions, *green lines* and *red lines* mark the reference-introns and test-introns, respectively; arrowheads mark 5’-end of introns, for which oligoprobes were designed. Positive and negative controls are highlighted by *green* and *red* fields, respectively. For clarity, RGB images of overlaid RNA signals are appended with grey scale images. Numbers of introns are indicated on the grey scale panels. Images are projections of short confocal stacks. Scale bars: *2 µm*.

## Notes

### Competing Interest Statement

The authors have declared no competing interest.

### Summary of Updates

No new experiments; only major revision of the text

